# Oxidants are dispensable for HIF1α stability in hypoxia

**DOI:** 10.1101/2020.05.05.079681

**Authors:** Amit Kumar, Manisha Vaish, Saravanan S. Karuppagounder, Irina Gazaryan, John W. Cave, Anatoly A. Starkov, Elizabeth T. Anderson, Sheng Zhang, John T. Pinto, Austin Rountree, Wang Wang, Ian R. Sweet, Rajiv R. Ratan

**Affiliations:** Burke Neurological Institute, White Plains, NY, 10605, USA; Brain and Mind Research Institute and Department of Neurology, Weill Medical College of Cornell University, New York, 10016, USA; Department of Neurology, Weill Medical College of Cornell University, New York, 10016, USA; Department of Pharmacological Sciences, Icahn School of Medicine at Mount Sinai, New York, 10029, USA; Department of Anatomy and Cell Biology, New York Medical College, Valhalla, NY, 10595, USA; Institute for Biotechnology, Cornell University, Ithaca, NY, 14853, USA; Department of Biochemistry and Molecular Biology, New York Medical College, Valhalla, NY, 10595, USA; Department of Medicine, University of Washington, Seattle, WA, 98108, USA; Department of Pain and Anesthesiology, University of Washington, Seattle, WA, 98195, USA

**Keywords:** Hypoxia, Mitochondria, Peroxide, Reactive oxygen species, HIF1α stability, S-glutathionylation, VHL, HIF PHDs, Oxygen, Ischaemia, Cancer, Stroke

## Abstract

Hypoxic adaptation mediated by HIF transcription factors has been shown to require mitochondria. Current models suggest that mitochondria regulate oxygen sensor (HIF prolyl hydroxylase) activity and HIF1α stability during hypoxia by either increasing mitochondrial peroxide as a second messenger or by serving as oxygen consumers that enhance the kinetics of cytoplasmic oxygen reduction. Here, we address the role of mitochondrial peroxide specifically in regulating HIF1α stability. We use state-of-the-art tools to evaluate the role of peroxide and other reactive oxygen species (ROS) in regulating HIF1α stability. We show that antioxidant enzymes are not homeostatically induced nor are peroxide levels increased in hypoxia. Forced expression of diverse antioxidant enzymes, all of which diminish peroxide, had disparate effects on HIF1α protein stability. Reduction of lipid peroxides by glutathione peroxidase-4 or superoxide by mitochondrial SOD failed to influence HIF1α protein stability. These data showed that mitochondrial, cytosolic and lipid ROS are dispensable for HIF1α stability and should affirm therapeutic efforts to activate the HIF pathway in disease states by HIF prolyl hydroxylase inhibition.

## Introduction

Over the past decade, our ability to monitor and manipulate reactive oxygen species (ROS) has grown enormously. These technological advances provide a novel view on how ROS interact with cells to modulate function. Specifically, ROS such as peroxide can act as cellular messengers. Messenger functions for ROS reflect their tight spatial control within cells. The tight spatial control of ROS has enabled their critical roles in growth factor signalling, inflammation, and regeneration (Hameed et al., 2015; Jain et al., 2013; Lei & Kazlauskas, 2014). Specific signalling roles for ROS are facilitated by the existence of motifs in proteins such as phosphatases that render these proteins specifically susceptible to redox modulation (Bae et al., 1997; Lee, Kwon, Kim, & Rhee, 1998; Salmeen et al., 2003).

An important area of biology where ROS signalling has been highly investigated but where no consensus has emerged is hypoxic adaptation. Seminal work from the Semenza, Kaelin and Ratcliffe groups has demonstrated that when oxygen tension falls below a critical threshold, that a family of enzymes dependent on oxygen, iron and 2-oxoglutarate known as the HIF Prolyl Hydroxylases (HIF PHDs) reduce their activity, leading to diminished hydroxylation of the alpha subunit of HIF transcription factors (Epstein et al., 2001; Ivan et al., 2002; Ivan et al., 2001; Jaakkola et al., 2001; Maxwell et al., 1999; Semenza & Wang, 1992; Wang, Jiang, Rue, & Semenza, 1995; Wang & Semenza, 1993). Diminished HIF1α hydroxylation reduces recruitment of a key E3 ubiquitin ligase, the von Hippel Lindau (VHL) protein, and this allows HIF1α to avoid proteasomal degradation. Stabilised HIF1α dimerises with its constitutively active partner to bind to hypoxia response elements in a coordinate gene cassette that leads to hypoxic adaptation at a cellular, local and systemic level (Jiang, Rue, Wang, Roe, & Semenza, 1996; Wang et al., 1995; Wood, Gleadle, Pugh, Hankinson, & Ratcliffe, 1996)). HIF transcription factors have been implicated in limiting damage to the kidney, heart and brain and in the progression of a host of cancers, as well as pulmonary fibrosis. Therefore, understanding the role of ROS in HIF1α-mediated adaptation has numerous clinical implications (Bryant et al., 2016; Conde et al., 2012; Semenza, 2012, 2014; Sheldon, Lee, Jiang, Knox, & Ferriero, 2014; Weidemann et al., 2008).

Previous studies have shown that the dose response of HIF1α stability that would progressively lower oxygen concentrations required other factors besides oxygen to regulate HIF PHD activity (Bell et al., 2007; Brunelle et al., 2005; Chandel et al., 2000; Mansfield et al., 2005). Using what were then state-of-the-art tools to monitor ROS, earlier researchers showed that ROS were increasing during hypoxia and that pharmacological tools that nullified this increase in ROS diminished HIF1α stability (Chandel et al., 2000). A compelling model emerged that hypoxia increases the flux of electrons via Rieske iron-sulphur (Fe-S) cluster proteins in complex III, and this leads to an increase in mitochondrial ROS generation via the ubiquinone binding site near the outer leaflet of the inner mitochondrial membrane (the Qo site) (Bell et al., 2007). Peroxide generated at this site could then diffuse through the outer mitochondrial membrane to inhibit HIF PHDs by either direct redox modulation of HIF PHDs (Bell et al., 2007) or by activation of established redox sensitive p38 MAP kinase signalling (Emerling et al., 2005).

Since its inception, the concept of a role for mitochondria in HIF signalling, as established by Chandel and Schumacker, has been validated (Agani, Pichiule, Chavez, & LaManna, 2000; Mansfield et al., 2005; Taylor, 2008). However, an alternate ROS-independent view of how mitochondria regulate HIF1α stability in hypoxia has also been advanced. Pharmacological inhibition of electron transport chain (ETC) complexes or genetic knockdown of Rieske Fe/S proteins or cytochrome c not only inhibit mitochondrial ROS production (Bell et al., 2007; Brunelle et al., 2005; Chandel et al., 2000; Mansfield et al., 2005), it also inhibits oxygen consumption. In this distinct scheme, inhibition of the ETC function would alter the kinetics of reduction of cytosolic oxygen levels. Indeed, several groups have provided data supporting the importance of respiratory chain-driven mitochondrial oxygen consumption in dictating cellular oxygen gradients. These gradients have been hypothesised to reduce oxygen concentrations to levels required to inhibit the enzymatic activity of oxygen-dependent HIF PHDs—the key upstream regulator of HIF1α stability (Chua et al., 2010; Doege, Heine, Jensen, Jelkmann, & Metzen, 2005; Hagen, Taylor, Lam, & Moncada, 2003).

In this manuscript, we leverage a host of complementary approaches that support the conclusion that peroxide is a dispensable mediator of regulated HIF1α stability in hypoxia. Unexpectedly, our results suggest that HIF1α stability in hypoxia is not oxidant-initiated.

## Results

### Antioxidant enzymes are not homeostatically induced in hypoxia

Increases in ROS that are sufficient for signalling or toxicity trigger homeostatic transcriptional increases in antioxidant enzymes (Christman, Storz, & Ames, 1989). To assess whether hypoxia results in similar homeostatic increases in antioxidant protein expression, we exposed human neuroblastoma (SH-SY5Y) cells to hypoxia for 8 h and measured protein expression levels of peroxisomal, cytosolic and mitochondrial antioxidant enzymes, including catalase, glutathione peroxidase-1 (GPX1), glutathione peroxidase-4 (GPX4), MnSOD and peroxiredoxin-3 (PRDX3) (Figure 1A). The 8 h time point was chosen to monitor homeostatic changes in antioxidant enzymes because this would be 6 h following observable HIF1α stability in hypoxia in SH-SY5Y cells, which provides adequate time for homeostatic increases to initiate transcriptional or post-transcriptional adaptations. At the 8 h time point, neither the protein level of the peroxisomal antioxidant catalase nor the antioxidants present in both cytosol and mitochondria, such as GPX1 and GPX4, showed any change in hypoxia with normalisation to actin (Figures 1B and 1C). However, since hypoxia induces HIF1α-dependent mitophagy (Aminova, Siddiq, & Ratan, 2008; Zhang et al., 2008), mitochondrial mass is decreased with increasing duration of hypoxia and includes decreases in mitochondrial DNA and proteins. Accordingly, we normalised distinct mitochondrially targeted antioxidant enzymes to the level of citrate synthase, a mitochondrial protein. When normalised to citrate synthase, the expression levels of mitochondrial antioxidants, such as MnSOD and PRDX3, also did not change during hypoxia (Figures 1D and 1E). We then endeavoured to establish whether these findings apply to non-transformed cells by studying the expression levels of antioxidant enzymes in hypoxia in primary neurons. Similar to the case in neuroblastoma cells, the antioxidant enzyme levels did not change in post-mitotic neurons during hypoxia (Figures 1F–1I). Taken together, these findings suggest that ROS levels do not increase during hypoxia.

**Figure 1.**
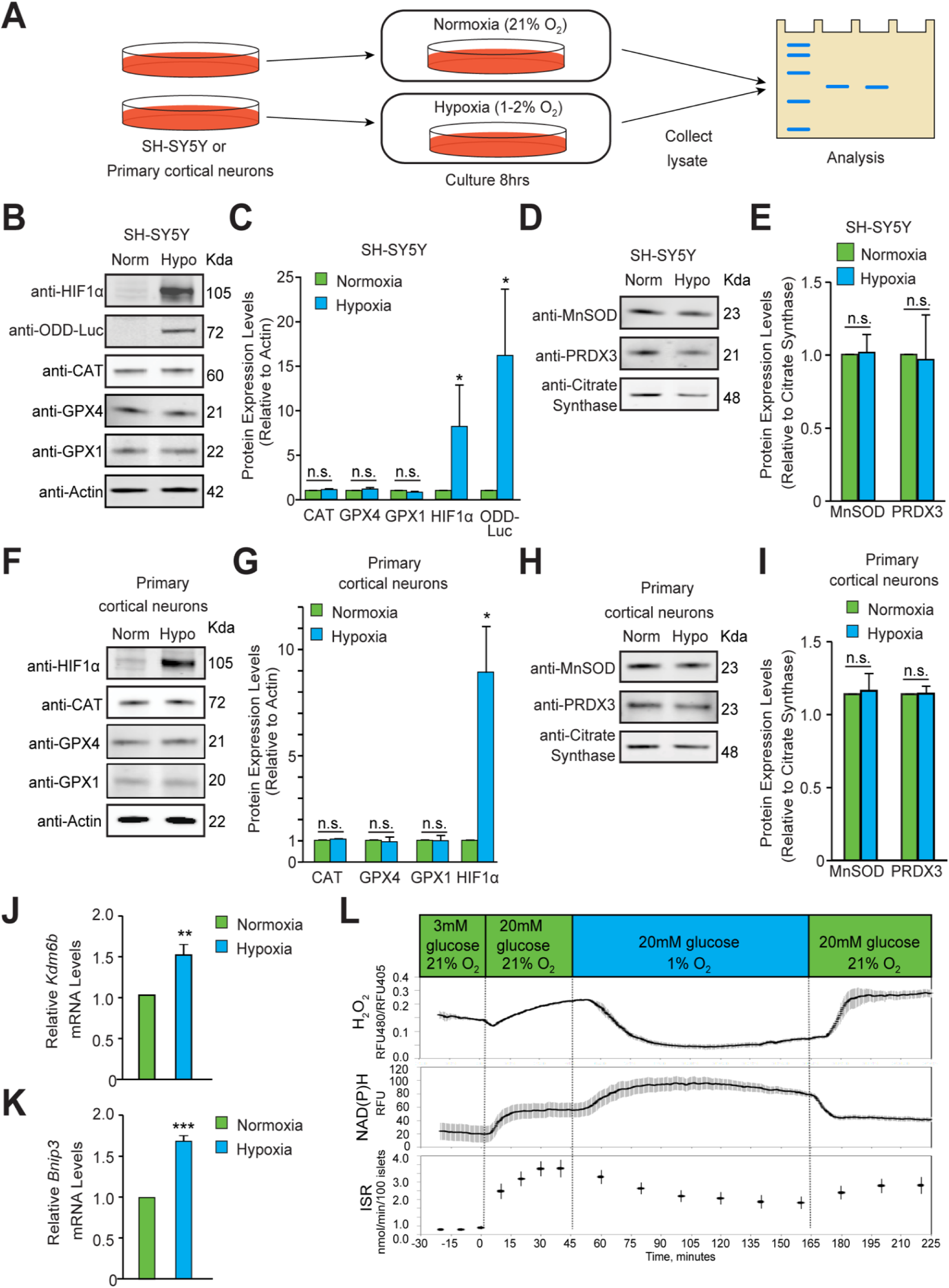
Hypoxia did not induce homeostatic increases in antioxidant enzymes or increase peroxide levels. **(A)** Experimental approach employed to examine changes in the protein levels of endogenous antioxidant enzymes in hypoxia. **(B-I)** Immunoblots of catalase (peroxisome), GPX1 and GPX4 (cytosol and mitochondria) or MnSOD and PRDX3 (mitochondria) in **(B-E)** SH-SY5Y oDD Luc. cells that stably overexpressed ODD (HIF **O**xygen **D**ependent **D**omain-luciferase fusion) or **(F-I)** PCNs exposed to normoxia or hypoxia for 8h. The protein levels of catalase, GPX4 and GPX1 were normalized to actin, while those of MnSOD and PRDX3 were normalized to the mitochondrial protein, citrate synthase. A monoclonal antibody to luciferase (indicated as anti-ODD-Luc. in the figure) was used to detect changes in ODD-luciferase protein levels in SH-SY5Y ODD-Luc. cells. **(J-K)** Pancreatic islets were exposed to hypoxia for 2h and then were lysed and processed for mRNA expression analysis of HIF1α target genes, *Kdm6b* and *Bnip3.* The densitometric data and gene expression data were pooled from three independent experiments an presented as mean ± SD. The densitometric data and gene expression data were statistically analysed using Student’s t test (B-K). (n.s.) indicates non-significant difference, (**) indicates p<0.01 and (***) indicates p<0.001 with respect to respective normoxia controls. **(L)** Hypoxia leads to a large decrease in H_2_O_2_ levels (top), increased NAD(P)H (middle), and decreased insulin secretion (bottom). Glucose stimulation by 20 mM glucose was added as reference, and oxygen levels were changed using an artificial gas equilibration device placed inline in the flow system. All three experiments were carried out separately, but using the same flow culture system.

### Peroxide levels do not increase during hypoxia

HyPer is a fusion protein composed of the peroxide-sensitive domain of the prokaryotic transcription factor, OxyR, and yellow fluorescent protein, which is a reporter for cellular peroxide (Belousov et al., 2006). This reporter is not only sensitive and specific, but its activity is also ratiometric, so this factors out any differences in fluorescence due to cell geometry, path length and reporter concentration. Prior studies have shown that enhanced pH buffering of the extracellular medium alleviates the putative effects of acidic pH during hypoxia on the reporter and that fluorescence ratios in cells can be calibrated to known peroxide concentrations (Neal et al., 2016).

We monitored changes in the level of hydrogen peroxide (H_2_O_2_) using real-time imaging of rat pancreatic islet cells under hypoxia with strong pH buffering. We used pancreatic islet cells, rather than neurons or neuron-like cells, for this initial analysis because mitochondrial peroxide in pancreatic islet cells increases in response in extracellular glucose levels, as measured using HyPer imaging. Elevated glucose can therefore be used as a positive control for mitochondrial peroxide increases. Elevated glucose also increases insulin release in beta islet cells, thereby enabling insulin release assays to establish cell viability in the absence of peroxide changes during hypoxia. Similar real-time measurements of cell function are not readily available in primary neurons or SH-SY5Y neuroblastoma cells.

Prior studies have also established that glucose-induced peroxide formation is derived from the mitochondria. Specifically, increased expression of mitochondrial catalase lowered the glucose-induced HyPer signals (catalase scavenges peroxide), as did reduced expression of mitochondrial SOD (MnSOD; the enzyme that converts superoxide to peroxide) (Neal et al., 2016). Finally, the signalling levels of peroxide measured by HyPer are 1/20th those of the peroxide levels required for toxicity (Neal et al., 2016), thereby confirming the sensitivity of the peroxide measurements using HyPer in this cell type. Before assaying peroxide in hypoxia, we verified that 2 h of hypoxia induced the established HIF1α target genes *Kdm6b* and *Bnip3* in islet cells in our flow culture system (Figures 1J and 1K) (Choudhry & Harris, 2018). This time point was selected to evaluate the role of peroxide in mediating the earliest changes in HIF1α stability in hypoxia because it occurs well before mitochondrial autophagy is induced (Figure 1).

Accordingly, based on the sensitivity and specificity of the reporter assay, we were confident that mitochondrial peroxide was measureable in islet cells should it increase during hypoxia. We measured H_2_O_2_, NAD(P)H and insulin secretion rates simultaneously as a function of glucose concentration for 2 h under hypoxic conditions (1% O_2_) (Figure 1L). Increasing the glucose concentration from 3 mM to 20 mM elicited the expected increases in peroxide, decreases in NADPH levels and insulin secretion rates under normoxia. Exposure of islet cells under the same glucose concentrations (20 mM) to 1% oxygen resulted in a decrease in H_2_O_2_ of more than 80%. This reduction occurred in the midst of increased NAD(P)H levels. NADP(H) likely increased due to its diminished utilisation by the mitochondrial electron transport chain. Glucose-stimulated insulin secretion decreased by about 50% in response to hypoxia, but it remained well above the unstimulated rates, indicating that the islets remained functional throughout the study. Indeed, restoring steady state O_2_ levels (20%) resulted in the expected increases in peroxide and insulin secretion, with concomitant decreases in NAD(P)H, indicating that hypoxia delivered under the conditions of our experiments is not toxic to islet cells (Figure 1L).

We then endeavoured to establish the generalisability of these findings to other cell types by measuring HyPer reporter fluorescence ratios in hypoxic in SH-SY5Y and Hep3B human hepatocellular carcinoma cells. Ratiometric imaging of both cell types showed no change in peroxide levels following 2 h of hypoxia, which was sufficient time to activate HIF1α-dependent gene expression. This absence of changes in peroxide levels during hypoxia could not be attributed to a lack of HyPer reporter responsiveness to peroxide in these cell types, since complex IV inhibition (KCN) (Figure S1C) or addition of exogenous peroxide (Figure S1G) following the hypoxic exposure led to the expected significant increases in reporter activity. Together with our antioxidant protein expression results, these data suggest that mitochondrial peroxide either decreases or is unchanged by hypoxia in primary (pancreatic beta islets) and transformed cell types (SH-SY5Y, Hep3B).

### HIF1α stabilisation is not oxidant initiated in hypoxia

We studied the regulation of HIF1α protein stabilisation in hypoxia directly by first establishing the sensitivity and dynamic range of HIF-luciferase reporter protein levels. The HIF-luciferase reporter contains the oxygen-dependent domain (ODD) of HIF1α fused to luciferase (ODD-luciferase). Prior studies from one of our groups have demonstrated that this reporter behaves like endogenous HIF but does not influence endogenous HIF activity (Karuppagounder et al., 2013; Smirnova et al., 2010). Treatment with deferoxamine (DFO), a canonical HIF PHD inhibitor, allowed dynamic monitoring of the HIF reporter protein levels across a wide range of DFO concentrations (Figure 2A and 2B). As an independent measure, we also assayed the activity of the HIF1α-luciferase reporter by measuring luciferase activity. These assays showed that the HIF1α-luciferase reporter measured by photometric luciferase activity possesses a high dynamic range and low coefficient of variation (Figure 2C). Indeed, the changes in HIF1α-luciferase activity measurements with increasing doses of DFO showed a strong correlation with the quantitative changes in HIF1α-luciferase reporter protein levels measured by quantitative immunoblotting (Figure 2D).

**Figure 2.**
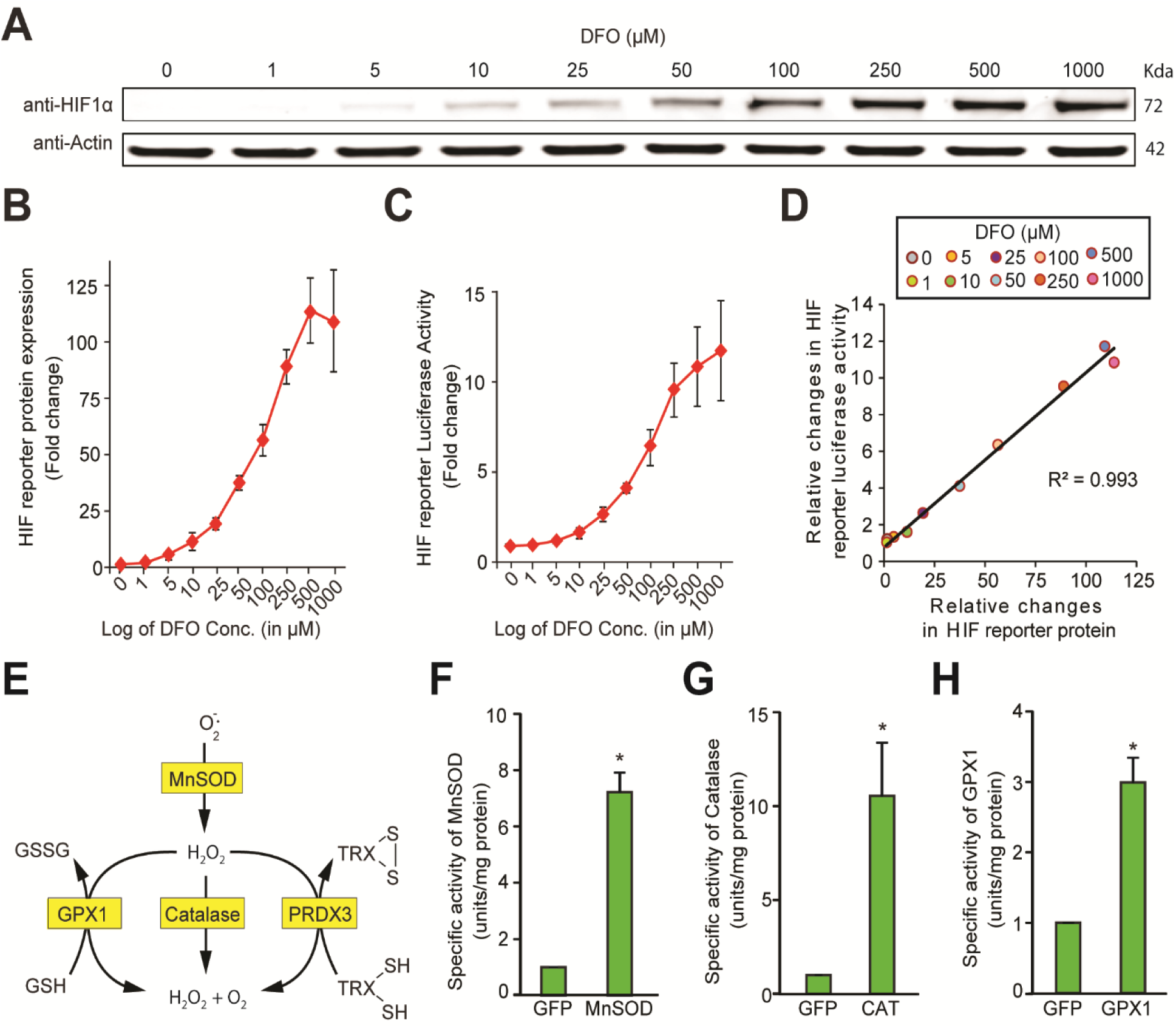
Validation of the sensitivity and the dynamic range of HIF1α protein and antioxidant enzyme activity measurements. **(A, B, C)** SH-SY5Y ODD-Luc. cells were treated with increasing concentrations of DFO (1 μM-1 mM) for 3 h and were, thereafter, processed for assessment of either changes in HIF reporter protein (i.e. ODD-Luc. protein) level via immunoblotting **(A, B)** or changes in HIF reporter luciferase activity (i.e. ODD-luciferase activity) with luminometry. **(C)**. **(D)** Correlation between relative changes in HIF reporter luciferase activity (i.e. ODD-luciferase activity) and relative changes in HIF reporter protein (i.e. ODD-Luc. protein). **(E)** A schematic diagram showing known mechanisms of H_2_O_2_ detoxification by peroxisomal, cytosolic and mitochondrial antioxidants**. (F, G, H)** SH-SY5Y cells were transduced with adenoviral constructs encoding MnSOD, catalase, GPX1 or a GFP control and incubated for 72h for steady-state expression. Cells were processed for measurement of specific activities of these enzymes. Data were pooled from three independent experiments in the form of mean ± SD. The statistical analyses were performed using Student’s t test. (*) indicates statistical difference of p<0.05 with respect to respective GFP control. “N” stands for normoxia and “H” stands for hypoxia.

Previous studies have shown that decreasing ROS by forced expression of individual antioxidant enzymes can decrease HIF1α protein levels (Brunelle et al., 2005; Chandel et al., 2000). We confirmed these findings with our HIF1α-luciferase reporter by forcing the expression of either catalase (a peroxide scavenger), GPX1 (a peroxide scavenger) or MnSOD (a superoxide scavenger and peroxide generator) (Figure 2E). SH-SY5Y cells expressing ODD-Luc were transduced with individual antioxidant enzymes encoded in distinct adenoviral constructs or an adenovirus encoding GFP only as a protein control. At 72 h following infection, GFP expression was observed in nearly 90% of SH-SY5Y cells (**Figures S2A and S2B**). Accordingly, the cells were tested for specific enzyme activities of MnSOD, catalase or GPX1. These studies showed a sevenfold, elevenfold and threefold increase in specific activity over GFP controls for MnSOD-, catalase-and GPX1-expressing cells, respectively (Figures 2F, 2G and 2H).

Upon verifying the increases in individual antioxidant enzyme activities (MnSOD, GPX1 and catalase) that can either increase (MnSOD) or decrease peroxide (GPX1, catalase), we then examined the effect of these manipulations on HIF1α stability. The mitochondrial peroxide model of HIF regulation predicts that MnSOD should increase HIF1α stability in hypoxia, whereas GPX1 and catalase should diminish HIF1α stability. In contrast to these predictions, we found that MnSOD had no effect on HIF1α reporter activity, whereas catalase increased reporter activity and GPX1 decreased it (Figure 3A). We further verified that our HIF1α luciferase reporter accurately reflected endogenous HIF1α levels by performing quantitative fluorescence immunoblotting. These assays showed that the changes in endogenous HIF1α protein are similar to those identified with the HIF1 a reporter (Figure 3B). From these findings, we concluded that HIF1α levels are not correlated with mitochondrial peroxide production.

**Figure 3.**
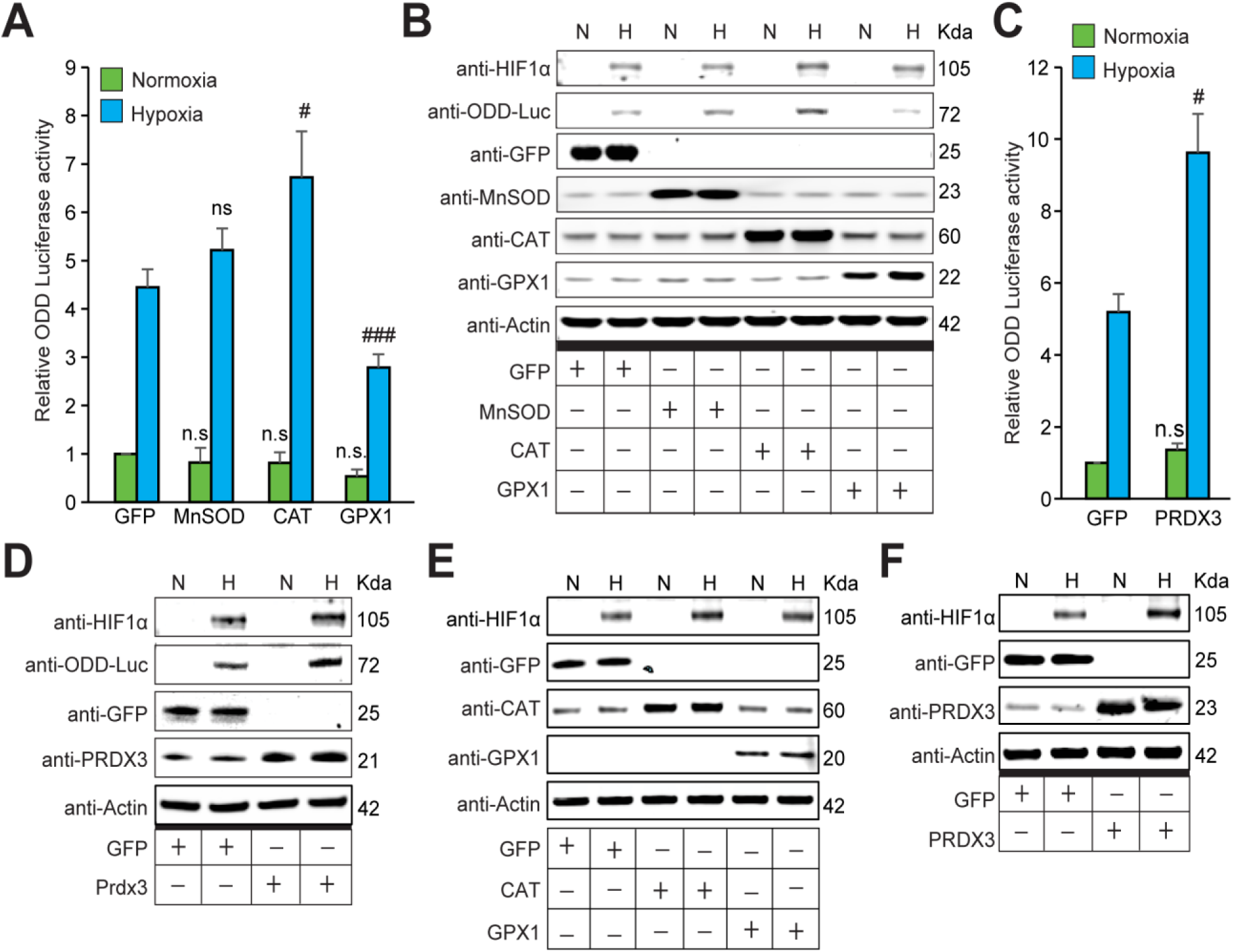
The stabilization of HIF1α is not oxidant-initiated in hypoxia (A, B, C, D) SH-SY5Y cells stably expressing ODD-luciferase were transduced with adenoviruses encoding distinct antioxidant enzymes for 72h and then exposed to normoxia or hypoxia in parallel and were either processed for luciferase activity assay (A measure of quantitative changes in ODD) **(A, B)** or immunoblot analysis (A measure of quantitative changes in protein levels of ODD and HIF1α) **(C, D)**. **(E, F)** Mouse primary neurons were transduced with adenoviral vectors encoding distinct antioxidant enzymes or GFP for 48h and then exposed to normoxia or hypoxia in parallel for 4h. Two way ANOVA with Bonferroni post-test was used for all comparisons. (n.s.) indicates non-significant difference with respect to GFP control under normoxia while (ns), (#) and (###) indicate non-significant difference and the statistical differences of p<0.05, and p<0.001, respectively, with respect to GFP control in hypoxia. All Western blot experiments were performed as three independent sets and a representative blot of each was shown in the figure.

Since catalase is a peroxisomal enzyme and GPX1 localises to the cytosol and mitochondria, our results could not formally exclude the possibility that GPX1 localisation to the mitochondria allowed it to reduce HIF1α levels, while the inability of catalase to penetrate this compartment did not allow it to reduce HIF1α levels. We addressed this possibility by forced expression of PRDX3, a member of the peroxiredoxin family of antioxidant enzymes that function as thioredoxin-dependent peroxide reductases in the mitochondria. Contrary to the GPX1 effects, forced expression of PRDX3 increased the levels of HIF1α during hypoxia, as measured by either HIF1α reporter activity (Figure 3C) or quantitative fluorescence immunoblotting (Figure 3D). We also confirmed these findings in primary neurons (Figures 3E and 3F), where adenoviral constructs also effectively increased expression of the antioxidant enzymes (**Figures S2C and S2D**). As a final test to confirm a reduction in oxidant production by GPX1, PRDX3 or catalase in SH-SY5Y cells, we measured the fluorescence of 5,6-carboxydichlorofluorescein (a non-selective redox sensitive reporter) using flow cytometry. These experiments confirmed the ability of GPX1, PRDX3 and catalase to reduce steady-state DCF oxidation, presumably resulting from oxidants generated physiologically (**Figures S3A and S3B**). However, no significant change was observed in the peroxide level in response to MnSOD expression, despite a significant increase in the MnSOD enzyme activity and protein level. This was unexpected and could reflect a compensatory activation of other antioxidants, such as GPX1, GPX4 or Prdx3, in response to increased MnSOD activity. We verified that DCF loading and the corresponding antioxidant effects were not different from normoxic or hypoxic cells, thereby arguing against the possibility that our redox reporter or the antioxidant enzymes are behaving differently in normoxia and hypoxia (**Figures S3D and S3E**).

Our findings did not exclude the possibility that antioxidants alter HIF1α protein levels by differential regulation of either *Hif* mRNA synthesis or *Hif* mRNA stability. Accordingly, we monitored *Hif1α* mRNA levels in the cells overexpressing catalase, GPX1 and PRDX3. Quantitative PCR revealed that *Hif* mRNA levels were not changed in a manner that would contradict the observed ROS-independent changes in HIF1α stability (Figures 4A and 4B). We confirmed that the changes observed in HIF1α protein stability are related to changes in its half-life by examining the stability of HIF1α protein in the presence of cycloheximide, which suppresses de novo protein synthesis in cells pre-treated either with or without MG132. As expected, we found that catalase and PRDX3, which increase HIF1α protein levels, also increased the HIF1α half-life (Figures 4C, 4D and 4E). By contrast, GPX1, which diminished HIF1α protein levels, decreased the HIF1α half-life (Figure 4D). Moreover, MG132 treatment significantly enhanced the HIF-1α half-life in all cases, showing that the antioxidant-led changes in hypoxic HIF1α stabilisation were a result of alterations in proteasomal degradation.

**Figure 4.**
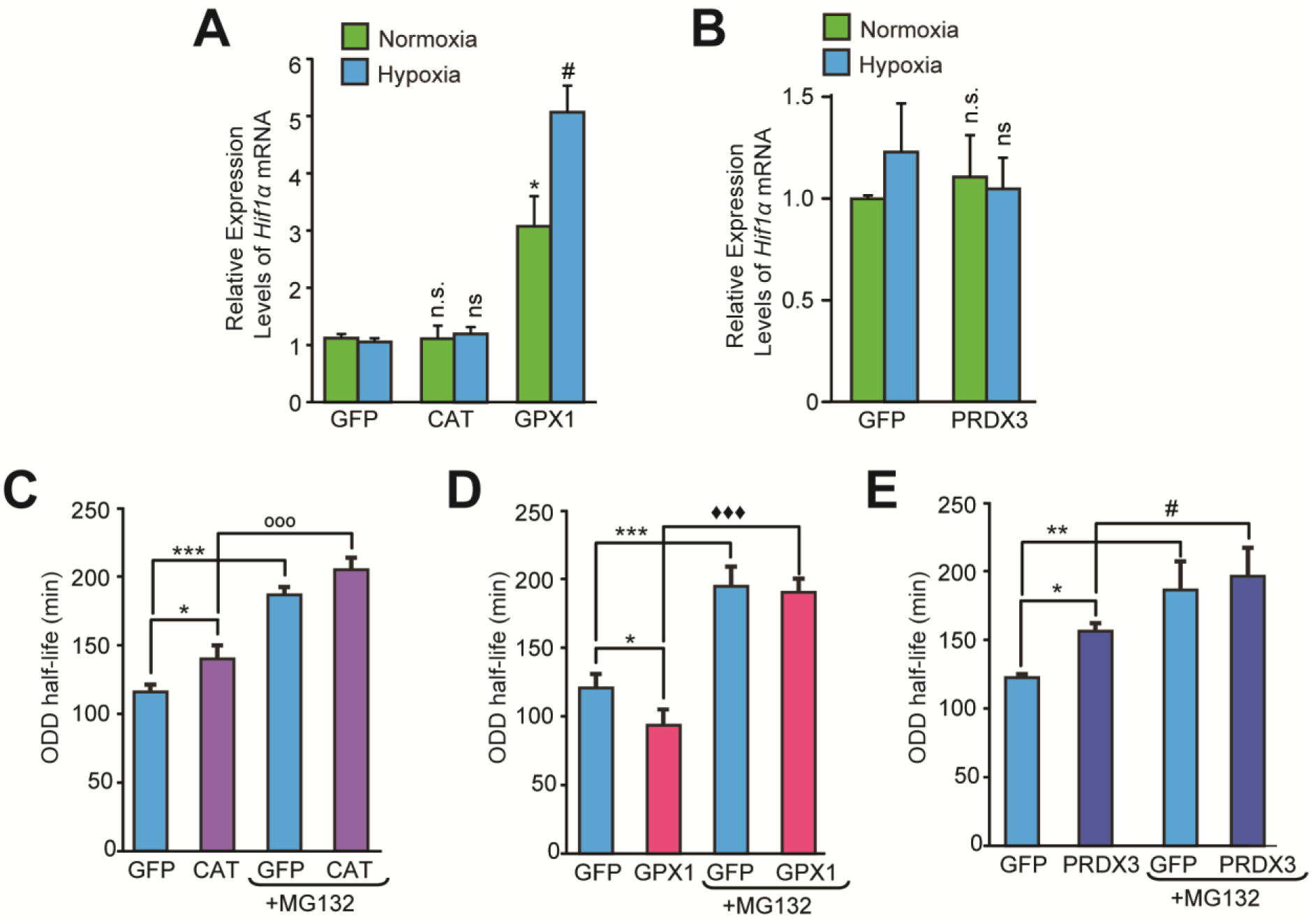
Divergent changes in HIF1α protein levels induced by antioxidant enzymes capable of scavenging peroxide cannot be attributed to differential changes in HIF1α mRNA synthesis or stability. **(A, B)** Relative changes in *Hif1α* mRNA in SH-SY5Y cells in response to forced expression of various antioxidant enzymes. Data were pooled from three independent experiments in the form of mean ± SD. One way ANOVA with Dunnett’s post-test was used for comparing cells expressing catalase or GPX1 with respect to cells expressing GFP and Student’s t test was used for comparing cells expressing PRDX3 with respect to GFP. (n.s.) indicates non-significant difference with respect to respective GFP controls under normoxia while (ns) and (#) indicate non-significant difference and the statistical difference of p<0.05, respectively, with respect to GFP control in hypoxia. **(C, D, E)** Changes in half-life of HIF1α in SH-SY5Y ODD-Luc cells expressing catalase or GPX1 or PRDX3 with respect to that of respective GFP controls in hypoxia. ODD Half-life was assessed by performing a pulse chase experiment by adding 35μM cycloheximide at every twenty minutes for a total of four hours using luciferase activity assay in SH-SY5Y cells expressing these antioxidants pre-treated either with or without 10 μM MG132. Data were pooled from three independent experiments in the form of mean ± SD. Two way ANOVA with Bonferroni’s post-test was used for statistical analysis. (*), (**), and (***) indicate statistical differences of p<0.05, p<0.01 and p<0.001 with respect to respective GFP controls in hypoxia. (°°°), (♦♦♦) and (#) represent statistical differences of p<0.001 w.r.t CAT, p<0.001 w.r.t GPX1 and p<0.05 w.r.t PRDX3, respectively.

We then examined whether our findings in primary neurons and neuroblastoma cells could be extended to non-neural cell types by examining the ability of antioxidant enzymes capable of modulating peroxide levels to modulate HIF1α stability in hypoxic Hep3B hepatocarcinoma cells and hypoxic HeLa cervical cancer cells. Forced expression of catalase, GPX1 or PRDX3 using adenoviral vectors significantly increased the protein levels of each of the antioxidant enzymes in Hep3B or HeLa cells (Figures S4 and S5). Similar to the effects in primary neurons or neuroblastoma cells, we did not observe a uniform reduction in HIF1α stability during hypoxia by forced expression of enzymes whose common activity is to reduce peroxide in these non-neural cell types (Figures S4 and S5). Taken together, our findings suggest that peroxide levels are uncoupled from HIF1α stability in both neural and non-neural cells.

### Is Factor Inhibiting HIF a target for ROS in hypoxia?

Previous studies have highlighted the role that Factor Inhibiting HIF (FIH) in the redox regulation of HIF1α. FIH inhibits HIF-dependent transcription by hydroxylating an asparagine in the C-terminal domain of the HIF1α protein that would normally interact with the co-activator CBP during hypoxia (Ema et al., 1999; Lando et al., 2002; Sang, Fang, Srinivas, Leshchinsky, & Caro, 2002). Suppression of FIH should, therefore, facilitate CBP recruitment and enhance HIF1 α target gene transcription. We determined whether a change in HIF-dependent transcription correlated with changes in HIF1α protein levels by examining *Enolase2* and *Bnip3* mRNA expression levels, as these are two established HIF1α target genes (Aminova et al., 2005; Poitz et al., 2014) in neuroblastoma SH-SY5Y cells and primary cortical neurons (PCNs). As expected, changes in *Enolase2* and *Bnip3* mRNA levels correlated with the HIF1α luciferase reporter levels (Figures 5A–5F). These results suggest that the differential regulation derives from differential effects of antioxidant enzymes on HIF1 α protein stability, rather than on FIH.

**Figure 5.**
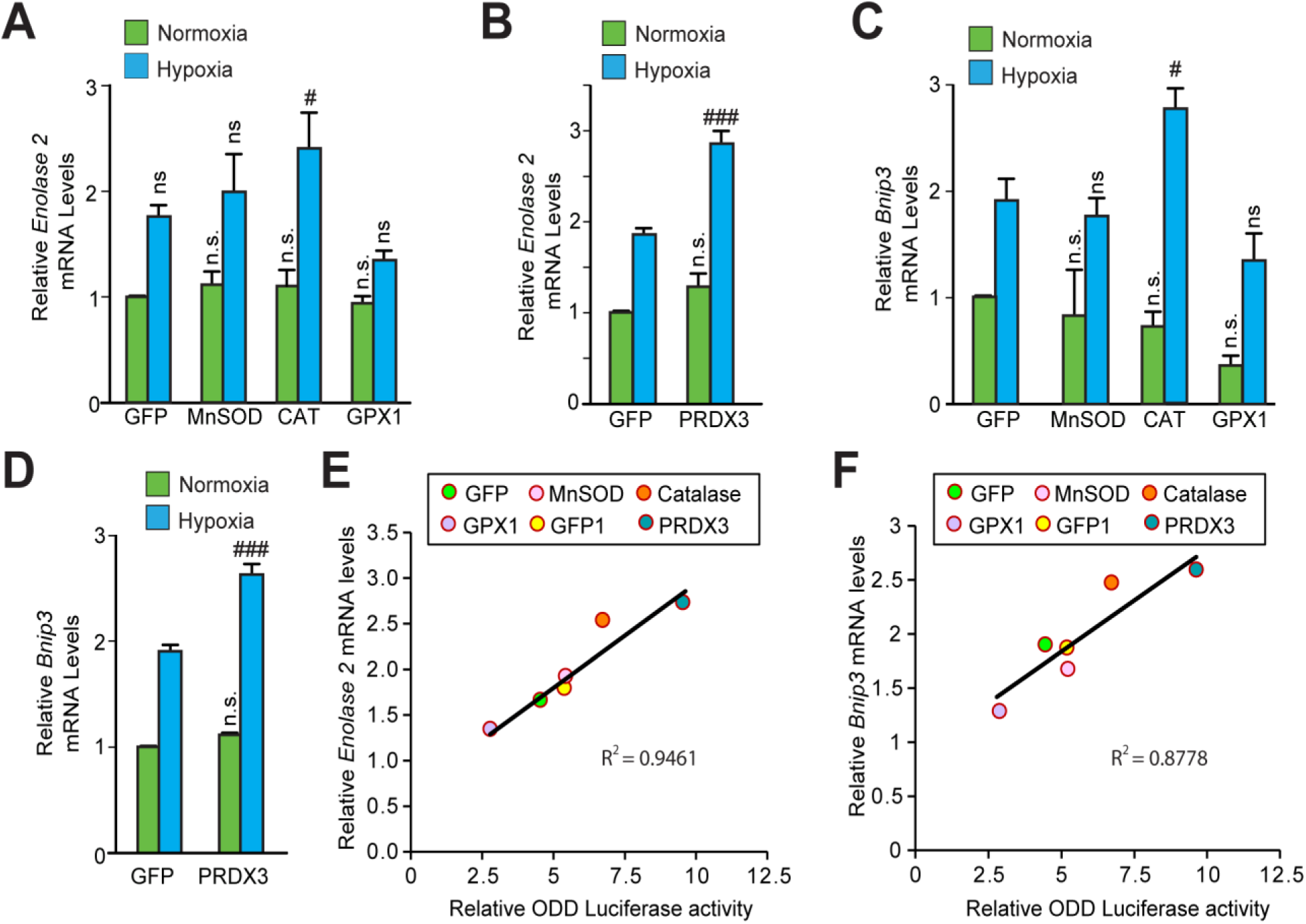
The transcriptional activity of HIF1α is not oxidant-initiated in hypoxia. Relative changes in mRNA levels of HlFla target genes **(A, B)** *Enolase2* and **(C, D)** *Bnip3* in SH-SY5Y cells expressing various antioxidant enzymes as compared to that of respective GFP controls. Data were pooled from three independent experiments in the form of mean ± SD. One way ANOVA with Dunnett’s post-test was used for comparing the statistical difference between cells expressing MnSOD, catalase or GPX1 with respect to that of GFP control and Student’s t test was used to compare the statistical difference between cells expressing PRDX3 with that of GFP control. (n.s.) indicates non-significant difference with respect to respective GFP controls under normoxia only while (ns), (#) and (###) indicate non-significant difference, and the statistical differences of p<0.05, and p<0.001, respectively, with respect to GFP control in hypoxia. **(E, F)** Correlation between relative changes in either *Enolase2* or *Bnip3* and relative ODD-luciferase activities.

### Neither Reactive Oxygen Species nor Reactive Lipid Species regulate HIF-1α stability in hypoxia

Recent compelling evidence has shown that reactive lipid species (RLS) are sufficient to drive HIF-dependent transcription via their effects on FIH inhibition without affecting HIF1α stability (Masson et al., 2012). Accordingly, we forced expression of GPX4, a selenoprotein that neutralises RLS (Figure 6A). We manipulated steady-state levels using the GPX4 protein fused to an optimised destabilisation domain (dd) from the prokaryotic dihydrofolate reductase gene. The dd domain destabilises GPX4 protein unless trimethoprim (TMP, 10 μM) is present (Figure 6B). TMP enhanced the GPX4 levels in neuroblastoma cells expressing ddGPX4, but increasing GPX4 levels had no effect on the hypoxia-induced HIF1α stability (Figures 6C–6D) or on HIF1α-dependent transcription (Figures 6E and 6F). We verified that GPX4 diminished RLS (Figure S3C) and neutralised the ferroptosis induced by glutamate, a form of cell death mediated by reactive lipid species that is abrogated by GPX4 (Figures 6G and 6H) (Tan, Wood, & Maher, 1998). Taken together, these findings argue against a central role for hydrogen peroxide or for lipid peroxides in mediating HIF1α stabilisation during hypoxia.

**Figure 6.**
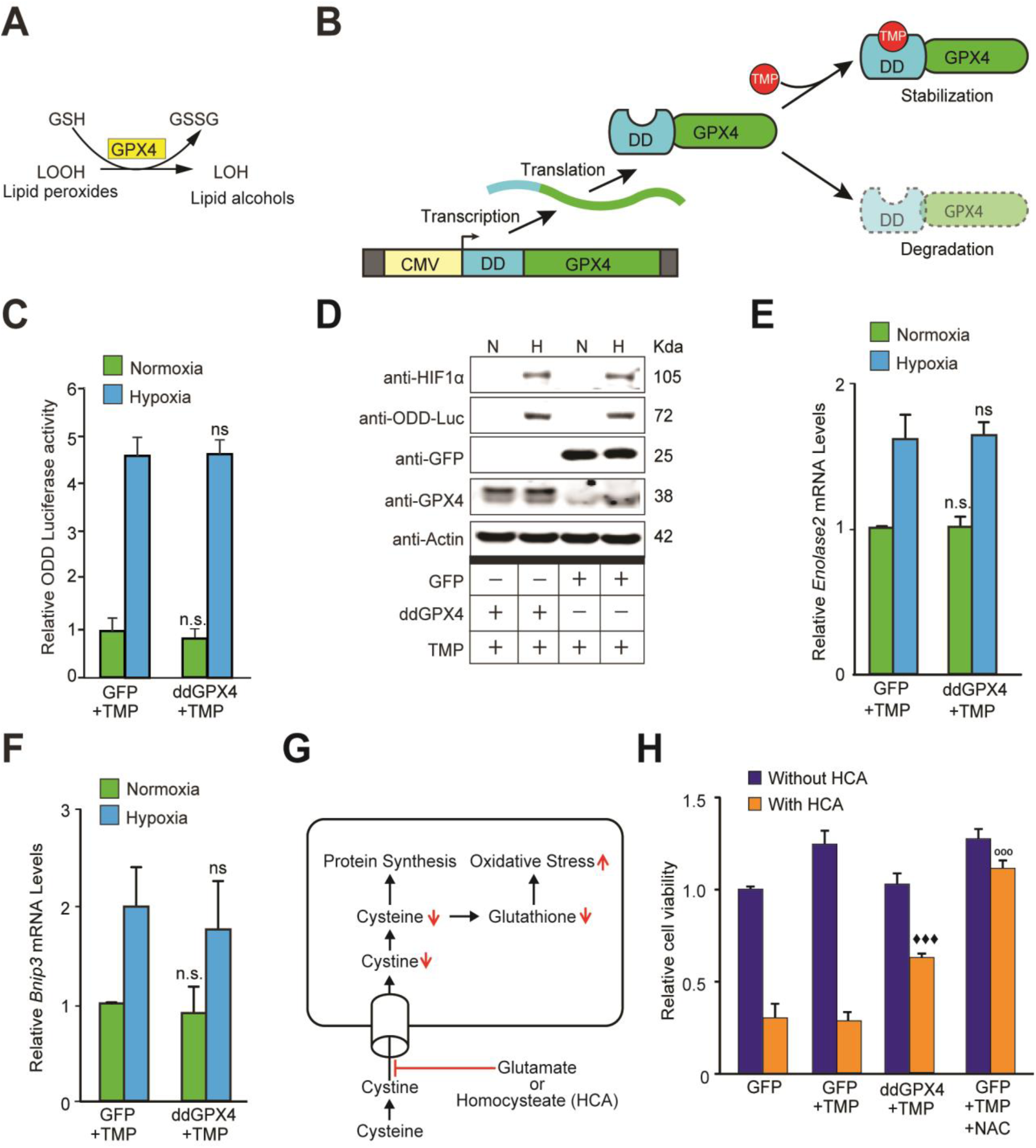
Reactive lipid species do not regulate HIF1α stabilization or its transcriptional activity in hypoxia. **(A)** A schematic diagram showing that GPX4 converts lipid hydroperoxides into lipid alcohols using GSH as a cofactor. **(B)** A schematic showing regulated protein expression of GPX4 fusion containing an optimized destabilization domain when exposed to the antibiotic, Trimethoprim (TMP). Reversible stability of GPX4 protein was conferred by fusing its coding sequence to a destabilization domain sequence (mutants of *E. Coli* dihydrofolate reductase). Accordingly, GPX4 protein possessing the destabilization domain is degraded resulting in low steady levels of GPX4. Trimethoprim binds to and neutralizes the destabilization domain stabilizing GPX4 protein in a dose dependent manner. **(C, D)** SH-SY5Y ODD-Luc cells were transduced with adenoviral vectors encoding a destabilized form of GPX4 (ddGPX4) or GFP and then exposed to normoxia/hypoxia and were either processed for luciferase activity assay (C) or immunoblotting (D). 10 μM TMP was added to ddGPX4 expressing cells after 60h of adenoviral incubation for 12h in order to achieve stabilized GPX4 expression. **(E, F)** Relative changes in mRNA levels of HIF1α target genes, *Enolase2* and *Bnip3* in SH-SY5Y cells. **(G)** A schematic diagram depicting the glutathione depletion model of oxidative stress. **(H)** PCNs were transduced with GFP/GPX4 for 24h and then treated with 5 mM HCA. Cells were simultaneously treated with 10 μM TMP. Then, cells were incubated for 24h to induce oxidative stress. Thereafter, viability of cells was measured via the MTT assay. 100 μM NAC was used as positive control. The final values were pooled as mean ± S.D. of three independent experiments. Two way ANOVA with Bonferroni’s post-test was used for statistical analyses. One way ANOVA with Dunnett’s post-test was used for comparing DCF fluorescence and viability. (n.s.), (*) and (***) indicate non-significant difference, and the statistical differences of p<0.05, and p<0.001, respectively, with respect to GFP control under normoxia while (ns), (#) and (###) indicate non-significant difference, and the statistical differences of p<0.05, and p<0.001, respectively, with respect to GFP control in hypoxia. (°°°) indicates statistical difference of p<0.001 with respect to GFP treated with HCA only while (♦♦♦) indicates statistical difference of p<0.001 with respect to GFP treated with both TMP and HCA.

## Discussion

Seminal studies have supported the notion that mitochondria are essential regulators of hypoxic adaptation and that they possibly act via their ability to generate peroxide (Agani et al., 2000; Chandel et al., 1998). In this paper, we show that the HIF1α stability mediated by HIF PHDs during hypoxia does not require peroxide. These data include our inability to detect an increase in peroxide during hypoxia (Figure 1L), the lack of homeostatic changes in antioxidant protein expression during hypoxia (Figures 1B–1I) and the failure of forced expression of antioxidant enzymes (catalase, GPX1, and PRDX3) that share a common ability to diminish cellular peroxide to influence HIF1α stability and transcription in the same direction (Figures 3 and 5). Our findings agree with prior studies that showed: 1) that HIF PHDs are not inhibited by exogenously added peroxide (Chua et al., 2010), and 2) that forced expression of an alternative oxidase which directly transfers electrons from coenzyme Q to oxygen to form water maintains HIF1α stability in hypoxia, despite reducing superoxide generation at Complex III (Chua et al., 2010).

Our results cannot be attributed to differences in mitochondrial ROS generation by transformed versus primary cells, or to differences in neuron-like versus non-neural cells, as SH-SY5Y neuroblastoma cells and primary cortical neurons showed similar effects, as did Hep3B hepatocellular carcinoma cells and HeLa cervical cancer cells. While we cannot exclude the possibility that culture conditions, such as serum lots, could reconcile our results with prior studies, in aggregate, the findings favour a role for mitochondria in modulating HIF1α stability via their effects as oxygen consumers rather than as peroxide second messenger generators.

### Peroxide scavengers have distinct effects on hypoxic HIF1α stability

Prior studies have shown that, in some cell types, ROS generation in hypoxia could be related to increased oxidant production or decreased defences (Naranjo-Suarez et al., 2012). We addressed the possibility that the imbalance of oxidants and antioxidants plays a regulatory role in mediating hypoxia signalling by the forced expression of distinct antioxidant enzymes known to either reduce (GPX1, catalase and PRDX3) or enhance (MnSOD) peroxide levels. Despite evidence for increased activity of the antioxidant enzymes studied using multiple experimental approaches, we found that HIF1α stability and transcription did not correlate with the effects on peroxide levels. Indeed, MnSOD had no effect on HIF1α stability, GPX1 diminished HIF1α stability and catalase and PRDX3 increased HIF1α stability in neuroblastoma cells and primary neurons (Figures 3A–3D). Similar uncoupling was observed in non-neural cell types as well (Figures S4 and S5). These results uncouple peroxide generation in the mitochondria from HIF1α stability via the HIF PHDs. One likely explanation is that the GPX1, catalase, and PRDX3 influenced HIF1α stability either via indirect but distinct effects on oxygen consumption or, alternatively, by differential but direct effects on proteins that influence HIF1α regulation.

### HIF1α stability under hypoxia is not regulated by reactive lipid species (RLS)

Recent studies have highlighted a potential role for RLS in regulating HIF transcription rather than HIF1α stability. Accordingly, we forced the expression of GPX4, a selenoprotein with known ability to neutralise reactive lipid species. Despite being active in combating RLS-mediated ferroptotic death in transformed cells (Figure 6H), GPX4 had no effect on HIF1α stability during hypoxia (Figures 6C and 6D) or on HIF-dependent transcription (Figures 6E and 6F). These data suggest that while some RLS (e.g. tert-butyl hydroperoxide) are sufficient to activate HIF transcription, they are not necessary for stabilisation of HIF1α or for driving HIF1α-dependent transcription during hypoxia.

Our collective data show that ROS or RLS are dispensable for HIF stability. The studies support a focus on HIF prolyl hydroxylase inhibition as a strategy for augmenting or increasing HIF activation in a host of disease states, including chronic kidney failure (to augment Epo), stroke and heart attack (Bryant et al., 2016; Conde et al., 2012; Semenza, 2012, 2014; Sheldon et al., 2014; Weidemann et al., 2008).

## Methods

### Cell lines and In Vitro Tissue Culture Studies

Immature primary cortical neurons were isolated from CD1 mice embryos (embryonic day 15 [E15]) as previously described (Ratan, Murphy, & Baraban, 1994) by following the protocol approved by IACUC at Weill Cornell Medicine. SH-SY5Y human neuroblastoma cells (purchased from ATCC) were cultured in DMEM/F-12 plus GlutaMAX medium added with 10% foetal bovine serum (Invitrogen) and 1% penicillin/streptomycin (Invitrogen). HeLa cells and Hep3B cells (purchased from ATCC) were cultured in EMEM medium added with 10% foetal bovine serum (Invitrogen) and 1% penicillin/streptomycin (Invitrogen).

Islets were harvested from Sprague-Dawley male rats (∼250g, Envigo, Huntingdon, Cambridgeshire, United Kingdom) anesthetised by an intraperitoneal injection of sodium pentobarbital (35 mg/230g rat). All procedures were approved by the University of Washington Institutional Animal Care and Use Committee. Islets were prepared and purified as previously described (Sweet et al., 2004) and then cultured at 37°C in RPMI Media 1640 (Gibco, Grand Island, NY) supplemented with 10% heat-inactivated foetal bovine serum (Atlanta Biologicals, Lawrenceville, GA) for specified times with the adenovirus coding the H_2_O_2_-sensitive dye HyPer.

### Adenoviral transduction and hypoxia/normoxia exposure

Adenoviral constructs of MnSOD, catalase GPX1 and respective GFP controls were purchased from the University of Iowa, Viral Vector Core Facility. Adenoviral constructs of ddGPX4 (GPX4 with destabilisation domain) and the respective GFP control were obtained from ViraQuest, Inc. (North Liberty, IA) and adenoviral constructs of PRDX3, GRX1 and the respective GFP controls were obtained from Vector Biolabs. For the GPX4 construct, we leveraged a novel technique recently developed by the Wandless group (Iwamoto, Bjorklund, Lundberg, Kirik, & Wandless, 2010) for deliberately regulating the level of expression of a protein of interest. An adenoviral construct of ddGPX4 had an *E. coli* dihydrofolate reductase (ecDHFR) mutant (called a degradation domain) fused to its CMV promoter which displays trimethoprim (TMP)-dependent stability. Because of the fusion of the degradation domain to the GPX4 promoter, GPX4 also displayed TMP-dependent stability. Without TMP, GPX4 was rapidly degraded completely through the proteasome, but increasing doses of TMP increased the GPX4 stability. Treatment of ddGPX4-expressing cells with 10μM TMP for 12 h stabilised ddGPX4 very well. SH-SY5Y, HeLa or Hep3B cells were transduced with different adenoviral constructs at 500 MOI (multiplicity of infection) and incubated for 72 h; primary immature cortical neurons (PCNs) were transduced with different adenoviral constructs at 100 MOI (Multiplicity of infection) for 48 h. The maximal expression of these constructs was determined by expressing the adenoviral construct of GFP at 500 MOI for 72 h in SH-SY5Y cells and 100 MOI for 48 h in PCNs on slides and staining them with GFP antibody (Abcam, ab6556). After 60 h of incubation, 10 μM TMP was added to ddGPX4 expressing cells and their respective control GFP expressing cells and the cells were incubated for a further 12 h to get stabilised GPX4 expression. Parallel sets without transduction of adenoviral constructs but probed with the same GFP antibody were used as negative controls in each in vitro model (Figure S2). Thereafter, one set was kept under normoxia (21% oxygen) and a parallel set under hypoxia (1% oxygen) for 4 h. The changes in endogenous antioxidants were studied by exposing SH-SY5Y cells and primary cortical neurons to normoxia/hypoxia for a relatively longer duration of 8 h to allow visualisation of a clearer difference under normoxic and hypoxic conditions.

Real-time changes in hydrogen peroxide during hypoxia were studied using an adenovirus containing the cytosolic H_2_O_2_ sensor, pHyPer-cyto vector (FP942, Evrogen, Moscow, Russia) (Belousov et al., 2006) generated by Vector Biolabs (Malvern, PA), as previously described (Karamanlidis et al., 2013). The H_2_O_2_ sensor was transduced into intact islets during incubation in RPMI media supplemented with 10% heat-inactivated foetal bovine serum and the adenoviruses at 100 MOI for 3 days at 37°C, as previously optimised (Neal et al., 2016). SH-SY5Y cells and Hep3B cells were also transduced with the H_2_O_2_ sensor at 100 MOI in a similar manner.

### Real-time epifluorescence imaging of intracellular H_2_O_2_

Real-time imaging experiments were carried out while islets were perifused using a commercially available temperature-controlled Bioptechs FCS2, which is a closed system, parallel plate flow chamber (Butler, PA) as previously described (Neal et al., 2016). After the islets were loaded into the perfusion chambers, the chamber was sealed and mounted onto the stage of a Nikon Eclipse TE-200 inverted microscope. KRB was pumped through the perifusion chamber at flow rates of 120 μL using a Masterflex L/S peristaltic pump (Cole-Parmer, Vernon Hills, IL). Use of an artificial gas exchanger positioned on the inflow side of the perifusion chamber enabled rapid changes in the concentrations of dissolved oxygen by switching the source gas tank between tanks containing 21% and 1% oxygen (balance 5% carbon dioxide and nitrogen)(Sweet, Cook, et al., 2002; Sweet, Khalil, et al., 2002). The HyPer signal was generated by dual fluorescence excitation with a xenon arc lamp (Lambda LS-1620, Sutter Instrument Company, Novato, CA) through either a 405/30nm or a 480/40 nm bandpass filter and detected at 510nm through a long-pass dichroic mirror with a cutoff below 500nm. The images were taken using a digital camera (Photometrics Cool Snap HQ2 CCD camera, Tucson, AZ) through a 40X Super Fluor Nikon objective (DIC H/N2). Data were expressed ratiometrically, where the excitation intensities at 480 nm were divided by those obtained during excitation at 405 nm. A similar procedure was used for real time monitoring of H_2_O_2_ in SH-SY5Y cells and Hep3B cells. We further confirmed the specificity of the HyPer signals by treating Hep3B cells with bacterial Streptolysin-O (which creates pores in the cell membranes) to selectively permeabilise the plasma membrane, followed by exposure to increasing concentrations of exogenous H_2_O_2_. Real time measurement of changes in the HyPer signals in response to exogenous addition of H_2_O_2_ confirmed the specificity of the HyPer signal with regard to H_2_O_2_.

### Real-time epifluorescence imaging of intracellular NAD(P)H

NAD(P)H autofluorescence was measured similarly to H_2_O_2_, except no dye loading was required and the excitation and emission wavelengths were 360 and 460 nm, respectively, as previously described (Gilbert, Jung, Reed, & Sweet, 2008). We calibrated the relative fluorescence units (RFU) at the end of experiments by measuring the steady state RFU in the presence of potassium cyanide (KCN) and, subsequently, FCCP. The normalised fluorescence of NAD(P)H was then calculated as follows:

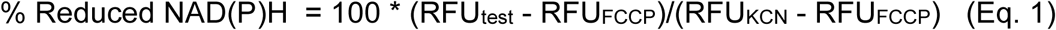

where RFU_FCCP_ and RFU_KCN_ equals the average of the final 10 time points during which each agent was present.

### Real-time measurement of insulin secretion rate

Outflow fractions from the flow system containing islets were collected in a fraction collector for subsequent measurement of insulin, as described previously (Sweet et al., 2004; Sweet, Cook, et al., 2002; Sweet, Khalil, et al., 2002). The insulin secretion rate was calculated as the flow rate (80 μL/min) times the insulin concentration in the outflow fractions, over the number of islets in the chamber, which was typically 50 (Sweet & Gilbert, 2006). Insulin was measured by radioimmunoassay (RI-13K, EMD Millipore, Darmstadt, Germany) as per the manufacturer’s instructions.

### ODD-luciferase activity assay

The ODD-luciferase construct with the pcDNA3.1 plasmid vector was constructed as previously described (Safran et al., 2006). The proline p402 and p564 present within the oxygen degradation domain (ODD) of HIF1α, when hydroxylated by HIF-PHDs, allow its binding to the VHL protein that targets it for proteasomal degradation. In this way, the stabilisation of ODD can be used as a marker of HIF1α stability (Safran et al., 2006; Smirnova et al., 2010). Because of the luciferase tagged with ODD, the increase in ODD stability leads to a proportional increase in the luciferase activity and this provides a very good way of measuring the HIF1α stability in a quantitative manner with a wide dynamic range. To this end, we used SH-SY5Y cells stably expressing ODD-luciferase. These cells were made by co-transfecting ODD-luciferase plasmid along with a puromycin resistance plasmid in SH-SY5Y cells, and stably transfected cells were positively selected in the presence of 4 μg/mL puromycin. Luciferase activity was measured by luciferase assay kit (Promega) using an LMaxII™ microplate luminometer (Molecular Devices). ODD-luciferase activity was normalised to the protein content of each well measured by Bio-Rad DC^TM^ protein assay kit.

### Gene expression study

Total RNA was prepared from SH-SY5Y cells using the Nucleospin RNA kit (Macherey-Nagel) and following their protocol. Real time PCRs were performed as a duplex reaction using FAM labelled *Enolase2* (Human - Hs00157360_m1), *Bnip3* (Human - Hs00969291_m1), and *Hif1α* (Human - Hs00153153_m1) gene expression assays (Thermo Fisher Scientific) and VIC labelled *human β actin* endogenous control probe (Human - 4326315E) or *RNA28S5* (Human - Hs03654441_s1) (Thermo Fisher Scientific) so that amplified mRNA can be normalised to *β actin* or *RNA28S5.* These experiments were performed using a 7500 Real-time PCR system (Applied Biosystems) using standard PCR protocols and amplification conditions.

We measure gene expression in pancreatic islets exposed to normoxia or hypoxia by first placing the islets in incubators containing either 21% or 1% oxygen for 2 h. The islets were then lysed and total RNA was purified using the RNeasy Mini Kit (Qiagen, Hilden Germany). *Bnip3, Kdm6b* and rat *Actin B* mRNA were measured by quantitative PCR using FAM labelled *Bnip3* (Rat - Rn00821446_g1), *Kdm6b* (Rat - Rn01471506_m1) gene expression assays and VIC labelled *rat β actin* endogenous control probe (Rat - 4352340E), all purchased from Thermo Fisher Scientific. These experiments were performed on an Mx3005P® Multiplex QPCR System (Stratagene, La Jolla, CA) with samples loaded in triplicate using ∼ 100ng of total RNA.

### Enzyme activity assays

For GPX1, catalase and MnSOD activity assays, cells in each sample expressing the relevant adenoviral constructs were collected, lysed and used for respective enzyme activity assays following the protocols of GPX1, catalase and SOD assay kits from Biovision. Total protein was measured using the Bio-Rad DC protein assay kit. The enzyme activity was normalised to the protein concentration for each sample.

### Cell viability assay

In order to test the functional activity of GPX4, immature primary cortical neurons (E15) were isolated from mouse embryos and plated at 10^6^ cells/mL in a 96-well plate. The next day, cells were transduced with GPX4 adenoviral constructs at 100 MOI. After a 24 h incubation, the cells were treated with the glutamate analogue, homocysteic acid (HCA) (5 mM), which inhibits the Xc^−^ transporter, thereby inhibiting cysteine uptake and leading to glutathione depletion and an increase in intracellular oxidative stress. Cells were also treated with 10 μM TMP at the same time to stabilise the GPX4 protein. The next day, cell viability was assessed by the MTT assay (Promega) to evaluate whether ddGPX4 is functionally active and shows its protective effect by decreasing oxidant levels under oxidative stress.

### ROS measurement through DCF flow cytometry

We measured changes in ROS levels using the molecular probe DCFDA (2’,7’-dichlorofluorescein diacetate). SH-SY5Y cells in normoxia as well as hypoxia were loaded with 20 μΜ DCFDA for 30 min. DCFDA diffuses through the cell membrane and is deacetylated by intracellular esterases to a non-fluorescent form, which is later oxidised by ROS into the highly fluorescent 2’,7’-dichlorofluorescein (DCF). Thereafter, parallel sets of cells in normoxia as well as hypoxia were treated with 5 mM H_2_O_2_ for 30 min as positive controls. After incubation, fluorescence was measured by flow cytometry at the wavelengths of excitation at 485 nm and emission at 535 nm. The production of ROS was measured as the mean fluorescence index multiplied by the respective cell counts and expressed as fold change with respect to the control.

### Immunoblotting

Protein extracts were prepared using 1% Triton buffer containing protease inhibitor, separated by SDS-PAGE, transferred onto nitrocellulose membranes and probed with antibodies against GFP (Cell Signaling Technology; 2555), MnSOD (Sigma-Aldrich; HPA001814), catalase (Sigma-Aldrich; C0979), GPX1 (Cell Signaling Technology; 3286S and Novus Biologicals; NBP1-33620), GPX4 (LSBio; LS B1596), PRDX3 (Novus Biologicals; NBP2-19777), Luciferase (Santa Cruz Biotechnology; sc-74548), HIF1α (Novus Biologicals; NB100-479) and citrate synthase (Cell Signaling Technology; 14309S).

## Quantification and Statistical Analysis

All experiments were performed as at least three independent sets, and data were displayed as means ± standard deviation (SD). Statistical significance was assessed in GraphPad Prism using either Student’s t test to compare values between two specific groups, one-way ANOVA followed by Dunnett’s post-hoc test/Tukey’s Post-hoc test to compare the values of more than two groups or two-way ANOVA followed by Bonferroni’s post-hoc test to compare the values of two groups under two different conditions at a given time. Statistical details for each figure can be found in their respective figure legends. The p value of 0.05 or less was considered statistically significant in all statistical analyses.

## Online Supplemental material

Fig.S1 provides additional evidence in SH-SY5Y cells or Hep3B cells that hypoxia does not increase peroxide levels. Fig. S2 shows a validation of the degree of expression of transduced transgenes encoded within adenoviral vectors in SH-SY5Y cell and primary cortical neurons (PCNs) using adenoviral particles encoding GFP. Fig. S3 shows the validation of the functional activity of antioxidants to reduce reactive oxygen species. Fig. S4 provides additional evidence in HELA cells that the stabilisation of HIF1α is not oxidant-initiated in hypoxia. Fig. S5 provides another additional evidence in Hep3B cells that the stabilisation of HIF1α is not oxidant-initiated in hypoxia.

## Acknowledgements

This work was supported by the National Institute of Health (Grant P01 AG14930-15A1, Project 1 to RRR), by a Dr. Miriam and Sheldon G. Adelson Medical Research Foundation grant to RRR, by a Goldsmith Fellowship to Amit Kumar for transition to independence, and by Diabetes Research Center Cell Function Analysis Core funding (P30 DK17047; University of Washington) to Ian Sweet. We also thank Sunghee Cho and Jiwon Yang for their help in data acquisition through flow cytometry. We also acknowledge the critical comments and suggestions from Drs. Ratcliffe, Schofield, Semenza, Silva and Ciechanover.

