## Supplemental data for "Oxidants are dispensable for HIF1α stability in hypoxia"

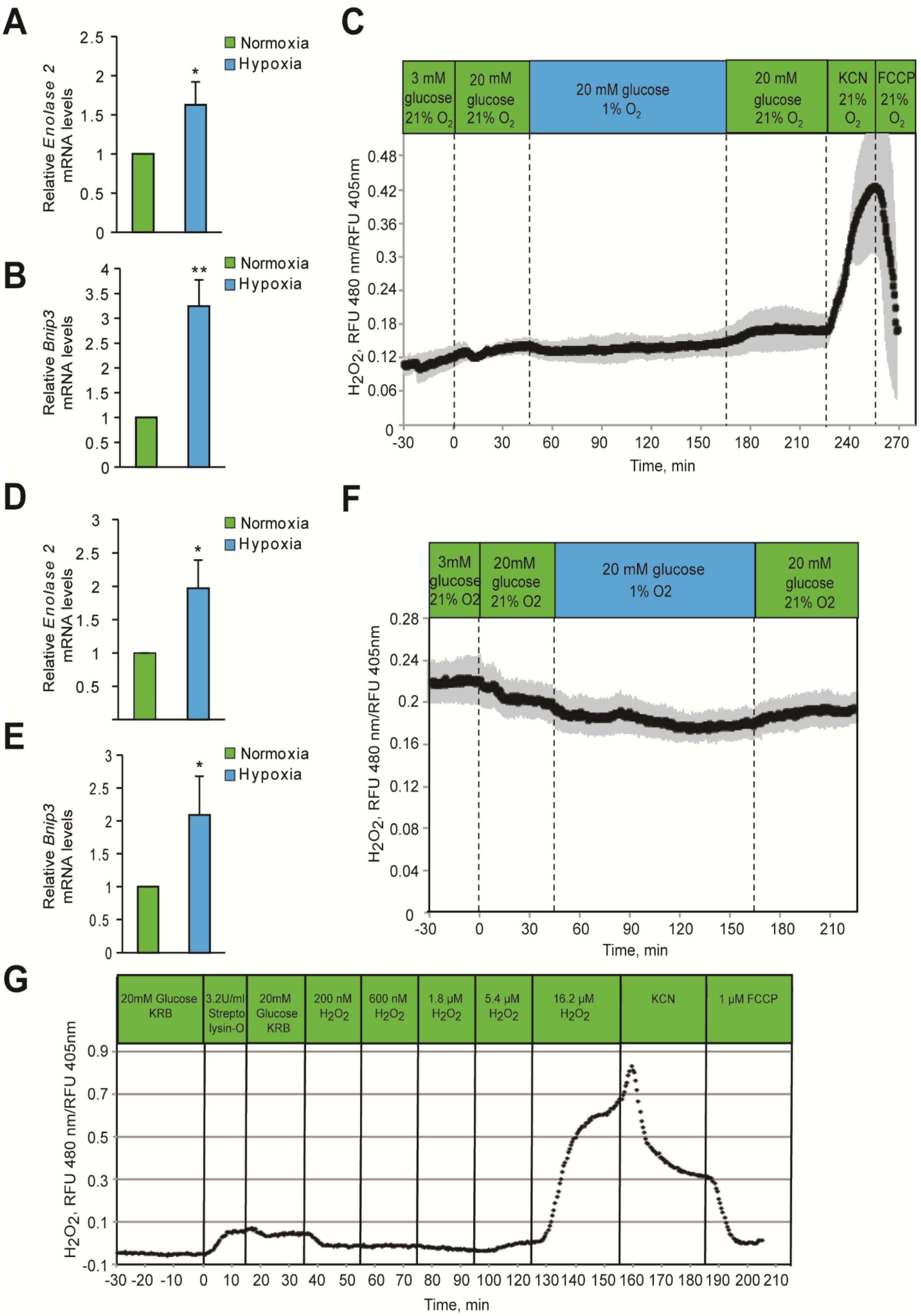
A
Relative Enolase 2 mRNA levels
Normoxia
Hypoxia
B
Relative Bnip3 mRNA levels
C
3 mM glucose 21% O2
20 mM glucose 21% O2
20 mM glucose 1% O2
20 mM glucose 21% O2
KCN 21% O2
FCCP 21% O2
H2O2, RFU 480 nm/RFU 405nm
Time, min
D
E
F
3mM glucose 21% O2
20mM glucose 21% O2
20 mM glucose 1% O2
20 mM glucose 21% O2
G
20mM Glucose KRB
3.2U/ml Strepto lysin-O
20mM Glucose KRB
200 nM H2O2
600 nM H2O2
1.8 μM H2O2
5.4 μM H2O2
16.2 μM H2O2
KCN
1 μM FCCP

**Figure S1. Hypoxia did not increase peroxide levels in SH-SY5Y cells or Hep3B cells.** SH-SY5Y cells **(A, B)** or Hep3B cells **(D, E)** were exposed to hypoxia for 2h and then were lysed and processed for mRNA expression analysis of HIF1α target genes, *Enolase 2* and *Bnip3*. The gene expression data was pooled from three independent experiments in the form of mean ± SD. The statistical analyses of gene expression data were done using Student's t test (J-K). (**) indicates $p<0.01$ and (***) indicates $p<0.001$ with respect to respective normoxia controls. **(C, F)** Hypoxia does not change $H_2O_2$ levels in either SH-SY5Y cells **(C)** or Hep3B cells **(F)**. Glucose stimulation by 20mM glucose was added as reference, and oxygen levels were changed by use of an artificial gas equilibration device placed inline in the flow system. **(G)** Hep3B cells, under normoxic conditions, were treated with bacterial Streptolysin-O to selectively permeabilize the plasma membrane followed by their exposure to increasing concentrations of $H_2O_2$. Real time measurement of $H_2O_2$ confirmed the specificity of Hyper signals with respect to $H_2O_2$. All three experiments were carried out separately, but using the same flow culture system.

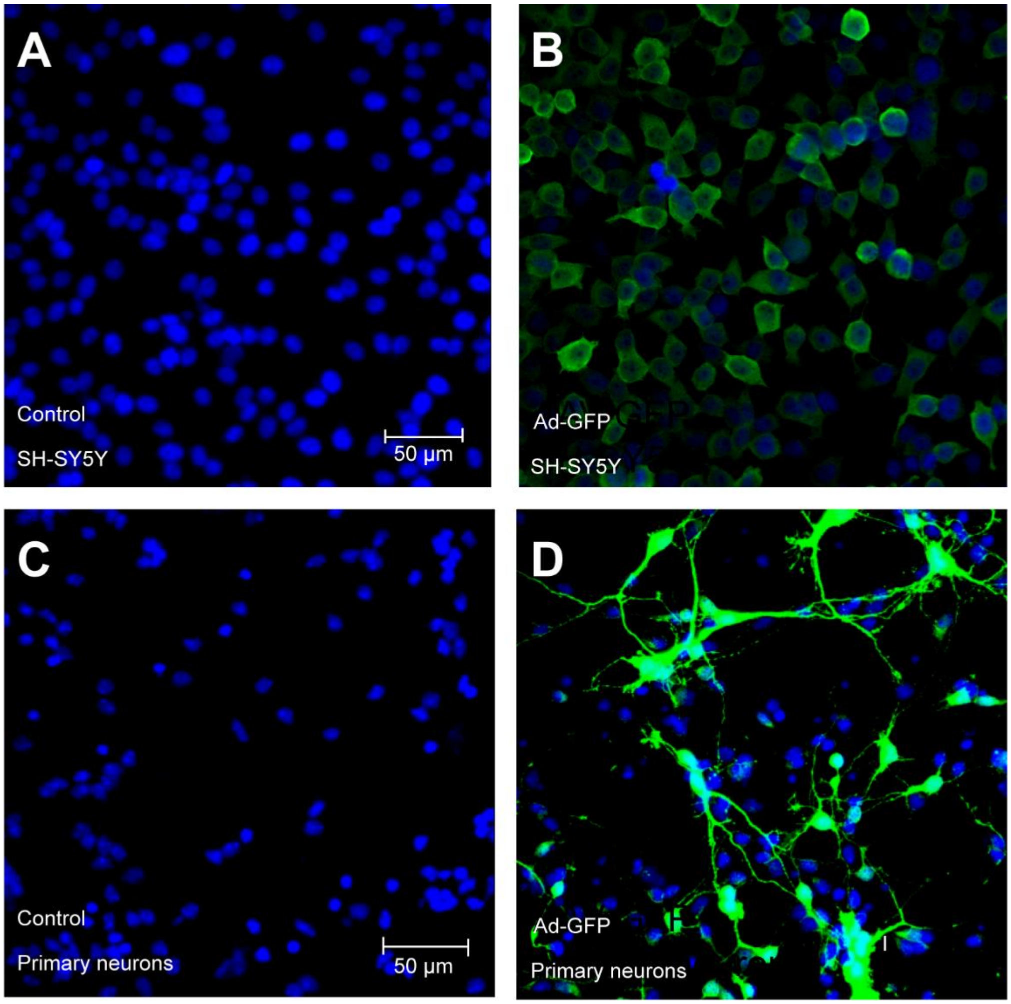


**Figure S2. Forced expression of transgenes encoded within adenoviral vectors in SH-SY5Y cells and primary cortical neuronal cultures (PCNs). (A-D)** Adenoviruses encoding GFP (Ad-GFP) were used to infect SH-SY5Y cells and PCNs, and expression was examined after 72h (A, B) and 48h **(C, D)**, respectively, to achieve steady-state levels of expression. Cells were immunostained using a GFP antibody to monitor the efficiency of overexpression. Parallel sets without adenoviral transduction were also monitored with GFP antibody and excluded non-specific staining. Experiments were performed as three independent sets and a representative picture of each is shown.

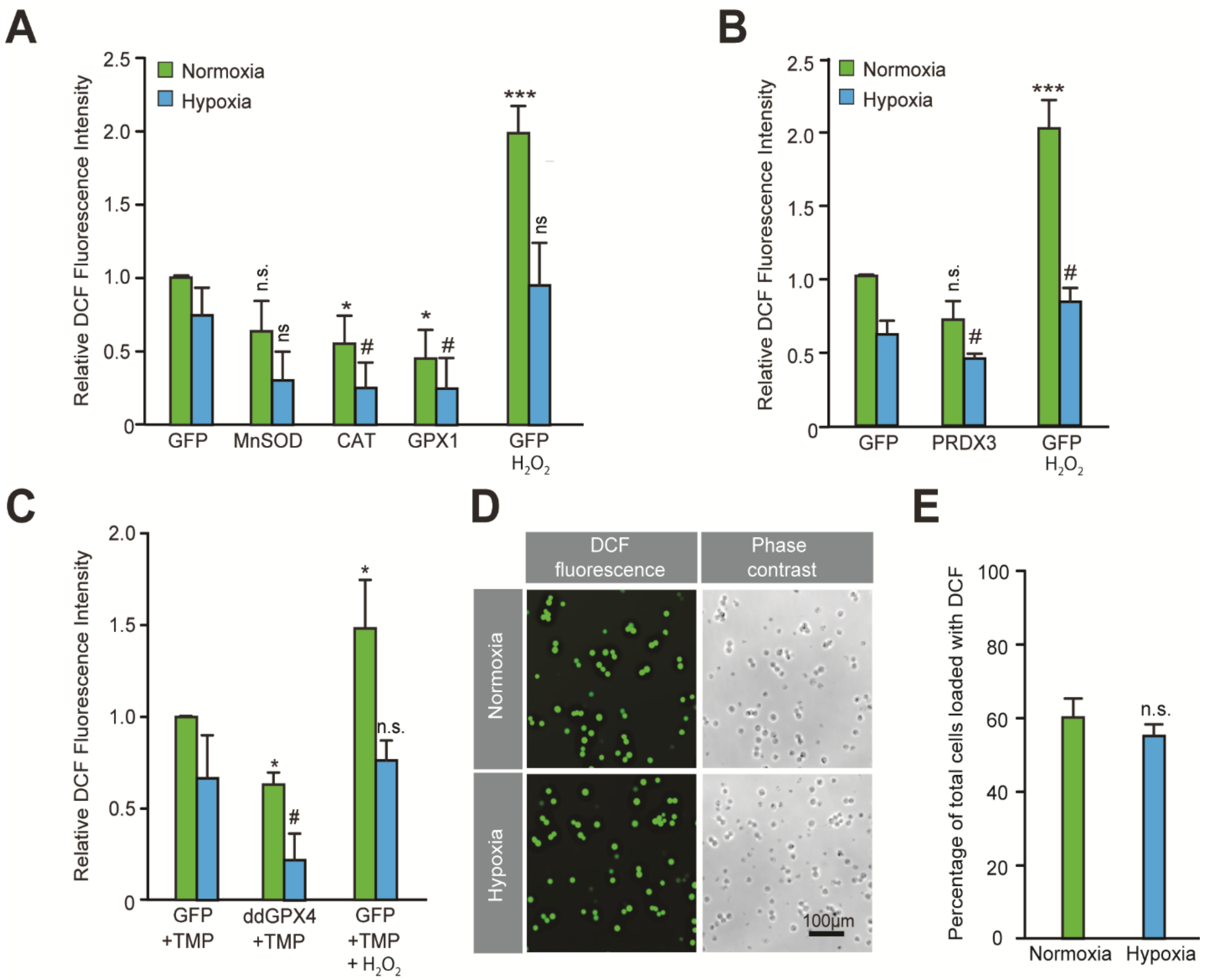


**Figure S3. Validation of the functional activity of antioxidants to reduce reactive oxygen species. (A-C)** The fluorescence of DCFDA loaded SH-SY5Y cells were measured through flow cytometry using standard fluorescein wavelengths. Fluorescence values were obtained by multiplying cell counts with mean fluorescence and later converted into relative fold change with respect to cells expressing GFP exposed to either normoxia or hypoxia. Parallel sets of cells in normoxia and hypoxia were treated with 5mM $H_2O_2$ for 30 min and were used as positive controls. The final values were pooled as mean ± S.D. of three independent experiments in all cases. One way ANOVA with Dunnett's post-test was used for all comparisons and a similar comparison was done in hypoxia separately. (n.s.), (*) and (***) indicate non-significant difference, and the statistical differences of $p<0.05$, and $p<0.001$, respectively, with respect to GFP control under normoxia while (ns) and (#) indicate non-significant difference and the statistical differences of $p<0.05$, respectively, with respect to GFP control in hypoxia. **(D, E)** DCF loading is equivalent in normoxic and hypoxic cells. SH-SY5Y cells were loaded with DCFDA for 30 min either in normoxia or hypoxia and then exposed to visible light equally to completely oxidize fluorescein. Illuminated cells incubated under normoxic or hypoxic conditions (from five independent sets) were counted manually to establish whether cells had equal DCFDA loading under both conditions. These values were pooled as mean ±

SD. The statistical analysis was done using Student's t test. (n.s.) indicates non-significant difference with respect to normoxia**.**

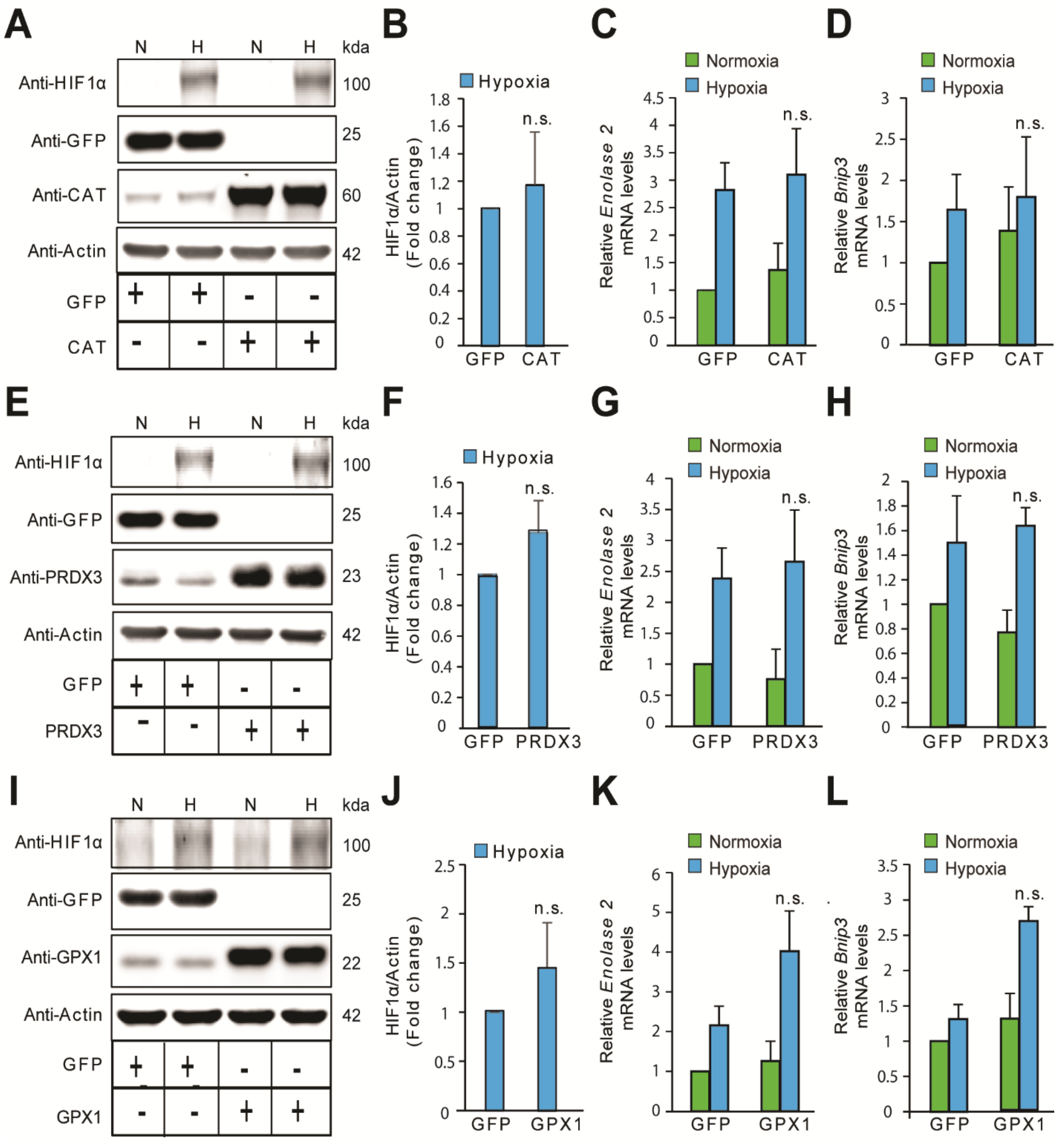


**Figure S4. The stabilization of HIF1α is not oxidant-initiated in hypoxia in HeLa cells.** HeLa cells were infected with adenoviruses encoding designated antioxidant enzymes and gene expression was assessed after 72h. Cells with forced expression of GFP or designated antioxidant enzymes were exposed to normoxia or hypoxia in parallel experiments and were processed for immunobloting of HIF1α or mRNA expression analysis of HIF1α target genes, *Enolase 2* and *Bnip3.* **(A, E, I)** Immunoblots showing changes in the protein level of HIF1α in response to catalase (CAT) overexpression (A), peroxiredoxin 3 (PRDX3) overexpression (E) or glutathione peroxidase 1 (GPX1) overexpression (I) under normoxia and hypoxia. (B, F, J) Densitometric analysis of changes in the protein level of HIF1α in hypoxia with designated antioxidant enyzmes. **(C, G, K)** Changes in mRNA level of HIF1α target gene, *Enolase 2* in respective overexpression conditions in normoxia and hypoxia. **(D, H, L)** Changes in mRNA level of HIF1α target gene, *Bnip3* in respective overexpression conditions in normoxia and hypoxia. The final values were pooled as mean ± S.D. of three independent experiments in all cases. Student's t test was used to compare the statistical difference between cells expressing antioxidants such as CAT, PRDX3 or GPX1 with that of respective GFP controls under hypoxia in densitometric analysis as well as gene expression analysis. (n.s.) indicates non-significant difference with respect to respective GFP controls under hypoxia. All western blot experiments were performed as three independent sets and a representative blot of each was shown in the figure.

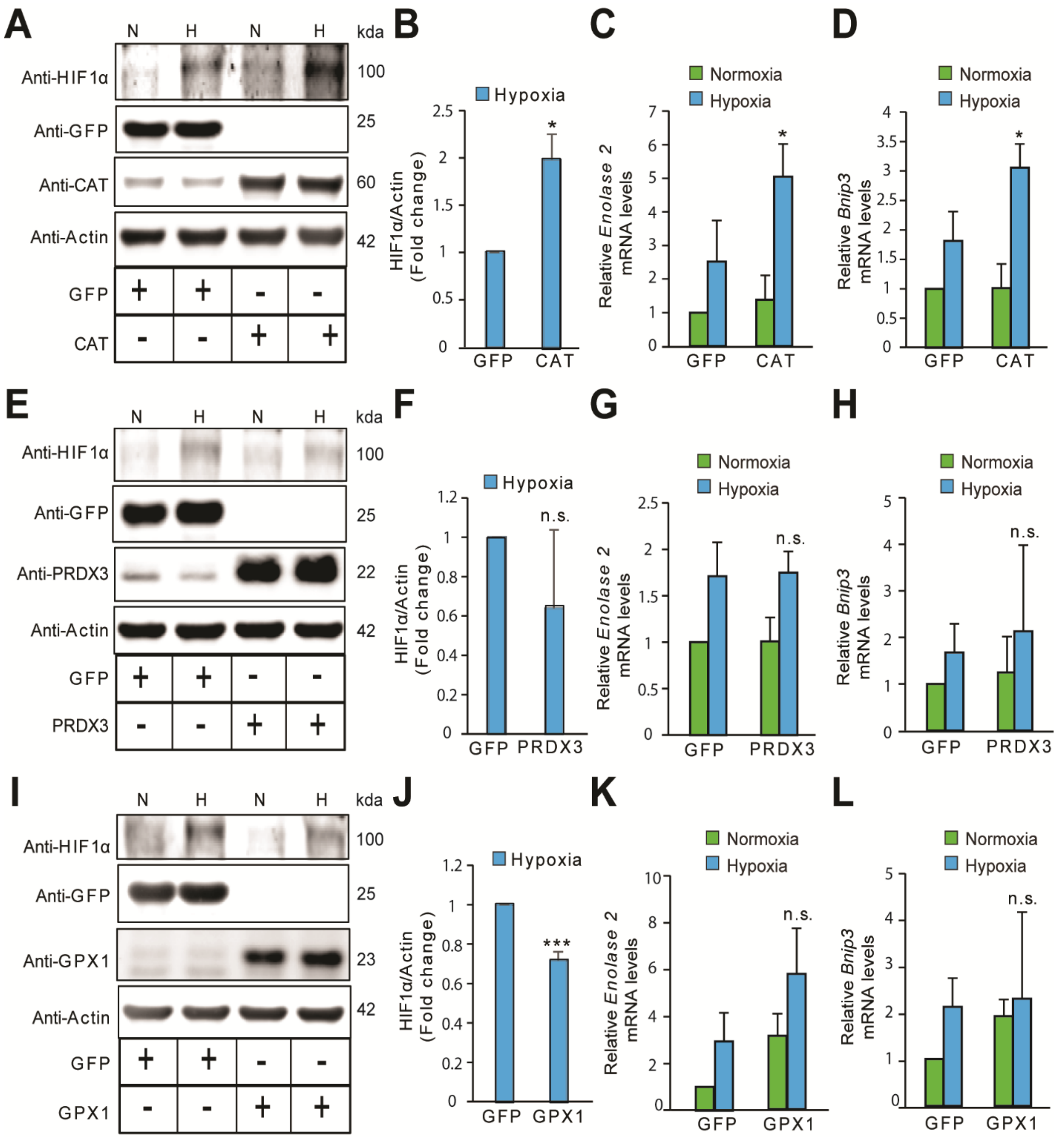


**Figure S5. The stabilization of HIF1α is not oxidant-initiated in hypoxia in Hep3B cells.** Hep3B cells were infected with adenoviruses encoding distinct antioxidant enzymes and transgenes reached steady state expression after 72h. Transduced cells were then exposed to normoxia or hypoxia in parallel experiments and were processed for immunoblotting to assess protein levels of HIF1α or mRNA analysis for assessing changes in levels of established HIF1α target genes, *Enolase 2* and *Bnip3.* **(A, E, I)** Immunoblots demonstrating changes in the protein level of HIF1α in response to either catalase (CAT) overexpression (A), peroxiredoxin 3 (PRDX3) overexpression, (E) or glutathione peroxidase 1 (GPX1) overexpression (I) under normoxia and hypoxia. **(B, F,**

**J)** Densitometric analysis of changes in the protein level of HIF1α under hypoxia in respective overexpression conditions. (C, G, K) Changes in mRNA level of HIF1α target gene, *Enolase 2* in respective overexpression conditions in normoxia and hypoxia. **(D, H, L)** Changes in mRNA level of HIF1α target gene, *Bnip3* in respective overexpression conditions in normoxia and hypoxia. The final values were pooled as mean ± S.D. of three independent experiments in all cases. Student's t test was used to compare the statistical difference between cells expressing antioxidants such as CAT, PRDX3 or GPX1 with that of respective GFP controls under hypoxia in densitometric analysis as well as gene expression analysis. (n.s.), (*) and (***) indicate non-significant differences, and the statistical differences of $p<0.05$, and $p<0.001$, respectively with respect to respective GFP controls under hypoxia. All immunoblot experiments were performed as three independent sets and a representative blot of each was shown in the figure.
